# Power-law growth models explain incidences and sizes of pancreatic cancer precursor lesions and confirm spatial genomic findings

**DOI:** 10.1101/2023.12.01.569633

**Authors:** Ashley L. Kiemen, Pei-Hsun Wu, Alicia M. Braxton, Toby C. Cornish, Ralph H. Hruban, Laura Wood, Denis Wirtz, David Zwicker

## Abstract

Pancreatic ductal adenocarcinoma is a rare but lethal cancer. Recent evidence reveals that pancreatic intraepithelial neoplasms (PanINs), the microscopic precursor lesions in the pancreatic ducts that can give rise to invasive pancreatic cancer, are significantly larger and more prevalent than previously believed. Better understanding of the growth law dynamics of PanINs may improve our ability to understand how a miniscule fraction of these lesions makes the transition to invasive cancer. Here, using artificial intelligence (AI)-based three-dimensional (3D) tissue mapping method, we measured the volumes of >1,000 PanIN and found that lesion size is distributed according to a power law with a fitted exponent of -1.7 over > 3 orders of magnitude. Our data also suggest that PanIN growth is not very sensitive to the pancreatic microenvironment or an individual’s age, family history, and lifestyle, and is rather shaped by general growth behavior. We analyze several models of PanIN growth and fit the predicted size distributions to the observed data. The best fitting models suggest that both intraductal spread of PanIN lesions and fusing of multiple lesions into large, highly branched structures drive PanIN growth patterns. This work lays the groundwork for future mathematical modeling efforts integrating PanIN incidence, morphology, genomic, and transcriptomic features to understand pancreas tumorigenesis, and demonstrates the utility of combining experimental measurement of human tissues with dynamic modeling for understanding cancer tumorigenesis.

## Introduction

Pancreatic ductal adenocarcinoma (PDAC), though rare, is predicted to be the second leading cause of cancer-related deaths in the United States by 2030.^1-3^ A major hurdle in confronting this aggressive disease is that there is no effective screening test for PDAC or its precursor lesions.^4^ As such, PDAC is often diagnosed late when distant metastases are present and few clinical options remain. Only 15% of patients present with localized disease at the time of diagnosis.^1^ Improved understanding of the early development of pancreatic cancer is a necessary first step to developing effective screening tools. The majority of PDACs are believed to develop from microscopic precursor lesions called pancreatic intraepithelial neoplasia (PanIN, Fig 1).^5^ Study of PanIN is uniquely complicated due to their small size: PanIN lesions cannot be seen through noninvasive diagnostic imaging such as computed tomography (CT), magnetic resonance imaging (MRI), and endoscopic ultrasound (EUS). PanINs can be studied in surgically resected tissues, and novel techniques for three-dimensional (3D) mapping of dense tissues at cellular resolution enable quantitative assessment of PanINs and the pancreatic microenvironment in histological images.^6-11^ Recent works utilizing a large cohort of 3D reconstructed human pancreata revealed that the pancreata of some individuals contain hundreds of PanIN lesions.^12,13^ This number contrasts with the relative rarity of PDAC and suggests that most PanIN lesions will never progress to cancer in a person’s lifetime. The mechanism governing this extensive PanIN initiation and growth in human tissues is poorly understood.

**Figure 1.**
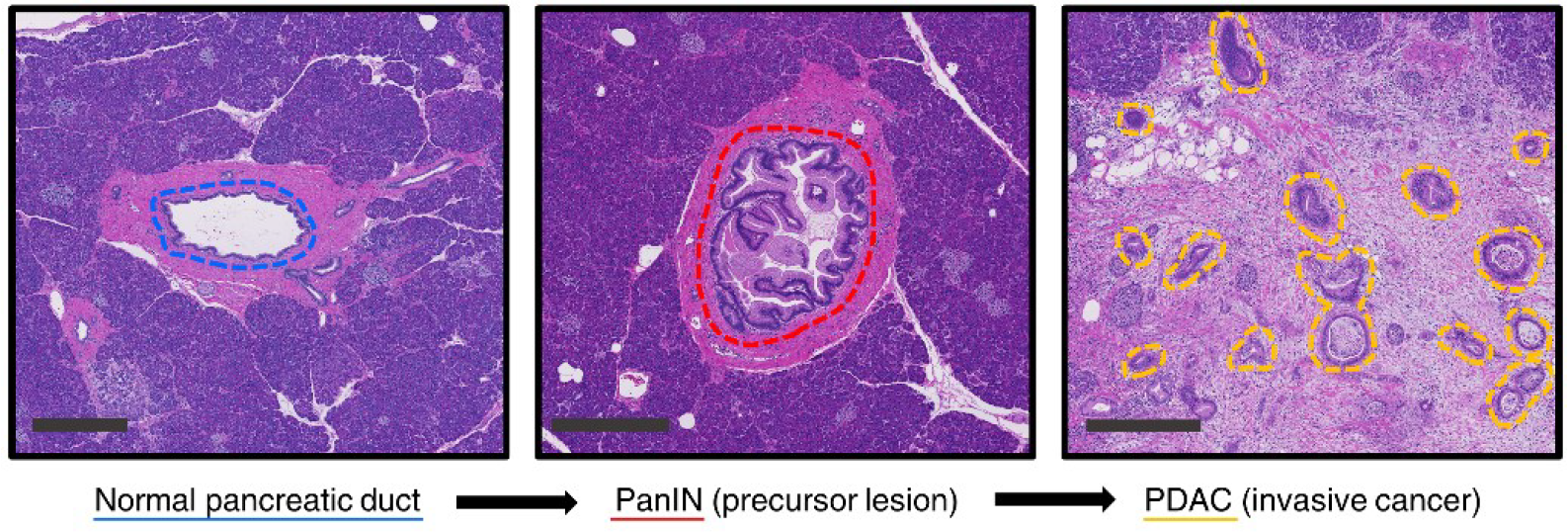
Pancreatic tumorigenesis as visualized in histological sections. Pancreatic ductal adenocarcinoma (PDAC) develops from histologically recognizable precursor lesions called pancreatic intraepithelial neoplasms (PanINs). Shown here are histological examples of (left) a histologically normal duct, (center) PanIN, and (right) invasive cancer. Scalebar = 0.5 mm.

The gold standard for understanding the true incidence and morphology of biological structures is direct measurement of 3D structure in human tissues. However, this approach has some limitations. Unlike a mouse model, where researchers maintain direct control over disease progression to pair structural metrics with temporal information, such control does not exist in study of human disease. Thus, while we can construct large cohorts containing structural information from hundreds of PanIN lesions, we do not know the ‘age’ of these precursors or possess information about the interrelation of adjacent lesions. Here, we utilize a cohort containing metrics from >1,000 PanIN lesions mapped in pancreatic tissues resected from 48 individuals to present potential growth dynamics of PanINs. Some of these samples contain spatially resolved DNA sequencing data describing the somatic mutations of spatially separate PanINs, providing additional information about their history.

Since PanIN growth cannot be observed directly, we use dynamic modeling to predict suitable growth laws by comparing the predicted size distributions to our experimental volumetric data.^14,15^ This approach allows us to identify fundamental processes contributing to growth. In particular, the spatially resolved genomics information suggests that intraductal spread of PanIN lesions, as well as multiple PanIN lesions fusing together to create large, highly branched structures might be important.^12^ In the following, we first analyze the experimental data in detail and then build successively more complex models to explain the data.

## Methods

### Experiments

#### Generation of 3D human pancreas tissue cohort

The 3D pancreas maps used here were previously described in a work mapping the prevalence and spatially resolved genomic properties of pancreatic cancer precursor lesions.^12^ Briefly, thick slabs of grossly normal human pancreas tissue were collected from 48 individuals who underwent surgical resection at the Johns Hopkins Hospital for pancreatic abnormalities including PDAC, well-differentiated pancreatic neuroendocrine tumors, metastatic disease of non-pancreatic origin, and non-malignant pathologies. Tissue was formalin-fixed, paraffin-embedded (FFPE), and serially sectioned at a thickness of 4µm. Every third section was stained with hematoxylin and eosin (H&E) and digitized at 20x magnification, for a lateral (xy) resolution of 0.5µm/pixel, and an axial (z) resolution of 12µm. CODA (Fig 2), a recently developed tool for 3D reconstruction of serially sectioned tissues,^16^ was used to register the serial images and segment nine pancreatic microanatomical structures on the serial H&E images at a reduced resolution of 2µm: PanIN, normal pancreatic ducts, pancreatic acini, islets of Langerhans, vasculature, nerves, fat, lymph nodes, and stroma to an accuracy of 96.6%.^12^ Resulting models were fully visualizable and quantifiable. Spatially distinct PanIN identified using CODA were validated through inspection of corresponding histology, and parameters including number of PanIN lesions per cm^3^ pancreas tissue, lesion size, cellularity, and aspect ratio were obtained.

**Figure 2.**
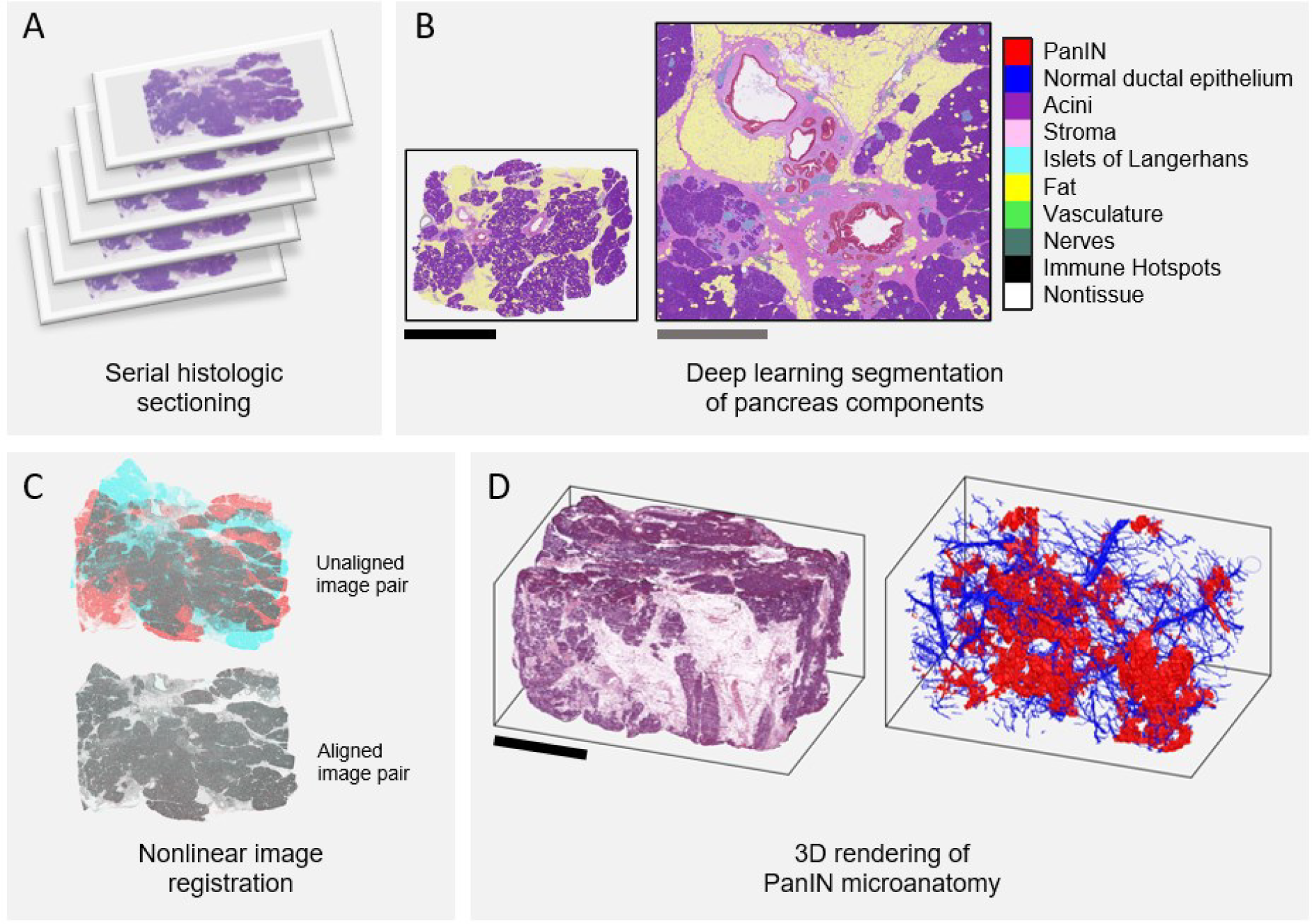
CODA 3D reconstruction of pancreatic microanatomy. (a) CODA starts with serial histological sectioning of formalin-fixed, paraffin embedded human pancreas samples. All, or a subset, of sections are stained with hematoxylin and eosin (H&E) and digitized. (b) A deep learning semantic segmentation algorithm was used to segment nine tissue components in the H&E images. (c) A nonlinear image registration algorithm was used to align the serial images into a digital volume. (d) Registered, segmented images were used to create visual and quantifiable maps of the pancreas microanatomy. Scalebars: black = 1 cm; gray = 2 mm.

### Power-law growth model

The power-law growth model given by Eq. (1) in the Results section with initial condition *V*(*t* = 0) = *V*_*min*_ results in the growth curve

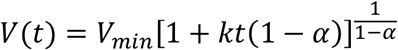

for *α* ≠ 1, and *V*(*t*) = *V*_*min*_*e*^*kt*^ for *α* = 1. The maximal PanIN size in a sample of age *T* is thus *V*_*max*_ = *V*(*T*), which is achieved when the PanIN is initiated at *t* = 0. Assuming a constant rate *j* of PanIN initiation, the distribution of PanIN sizes can be expressed by the complementary cumulative distribution function (CCDF) after duration *T*, which reads

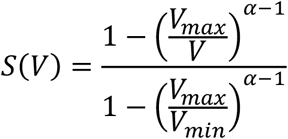

for *α* ≠ 1, and *S*(*V*) = *ln*(*V*_*max*_ /*V*)/*ln*(*V*_*max*_ /*V*_*min*_) for *α* = 1. These expressions do not depend on the initiation rate *j* since the distribution is normalized, *S*(*V*_*min*_) = 1. Moreover, inverting the growth curve *V*(*t*) allows us to determine the duration it takes for a PanIN to grow to the observed volume *V*, so we can predict when a PanIN must have been initiated for given *k* and *α*. Choosing the minimal plausible value for *k* (such that PanINs must have originated after *t* = 0) as well as pooling observed data by years and weighing them with the inverse sample volume, we predict the number *N*_*i*_ of PanINs that were initiated in year *i* per unit volume of the sample. This discretized data is smoothed with a Gaussian filter of width 5 *y* to generate Fig. 4F.

### Simulating extended models

We simulate the seeding model by explicitly propagating forward in time a collection of PanIN sizes {*V*_*i*_}. For each step, we first use Eq. (2) (Results section) to determine the average waiting time *Δt* until a new PanIN is initiated, and then grow all PanINs for this duration according to the power-law growth curve given above. We quantify the resulting distribution *S*^*pred*^(*V*) from the final volumes after time *T*.

In contrast, we simulate the merging model using a fixed time step *Δt* = 0.01 *y*. During each step, we first grow all PanINs according to the power-law growth given by Eq. (1), then randomly choose *M* of the *N*_*pairs*_ = *N*(*N* − 1)/2 possible PanIN pairs for an attempted merge, and finally initiate new PanIN with the constant initiation rate *j*_0_. The merge is performed stochastically, i.e., when *P*_*ij*_ > *ξ*, where *P*_*ij*_ = *Δt* · *N*_*pairs*_ *K*(*V*_*i*_, *V*_*j*_)/*M* with *K*(*V*_1_, *V*_2_) given by Eq. (3), and *ξ* is a random number chosen uniformly between 0 and 1. Here, *M* is a control parameter, which is chosen minimally while still obeying *P*_*ij*_ < 1. While initiation is still implemented deterministically, merging is done stochastically, so we obtain the respective distribution *S*^*pred*^(*V*) from an average of 8 independent runs. In all cases, we run simulations for *T* = 65 *y*, the median age of the patients analyzed. The volume *V*_*S*_ of the model sample does not affect results and we chose *V*_*S*_ = 100 *cm*^3^ to get adequate statistics.

To fit the predictions of these models to the observed data *S*^*obs*^(*V*), we minimize the logarithmically scaled mean squared deviation

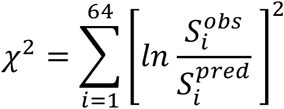

where the CCDFs *S*(*V*) are evaluated in 64 logarithmically distributed intervals between *V*_*min*_ and the maximally observed volume, so *S*_*i*_ = *P*(*V* > *V*_*i*_) gives the fraction of PanIN with a volume above *V*_*i*_. We minimize *χ*^2^ by adapting the model parameters using the Differential evolution algorithm^17^ over 8 independent repetitions, each with 2048 steps.

## Results

### PanIN sizes exhibit a broad distribution

The CODA methodology was successfully used to map PanIN lesions in human pancreas tissues (Fig 2). From each sample, we compiled patient demographic information along with number and size of PanIN lesions per 3D reconstructed surgically resected pancreas sample. Using these data, a range of PanIN sizes and morphologies was found (Fig 3A). A total of 48 thick slabs of human pancreas tissue were assessed (Fig 3B). The mean sample volume was 1.83 cm^3^ (median: 1.87 cm^3^, range: 0.31 – 3.62 cm^3^). Samples contained an average 21.8 spatially separate PanIN lesions (median: 18.5, range: 4 – 92). PanIN volumes were highly variable within this cohort. The smallest PanIN was 9 × 10^−5^ mm^3^, occupying part of a small, intercalated duct, and the largest PanIN was 24.7 mm^3^, occupying most of the pancreatic ductal system of the sampled region. The average PanIN volume was 0.27 mm^3^ (median 0.01 mm^3^). PanIN structure was similarly highly variable, with small PanIN lesions occupying short regions of single duct branches, and the larger PanIN lesions appearing highly branched, with extension in the pancreatic ducts and into surrounding acinar lobules. Figure 3C displays PanIN densities per sample, calculated as number of PanIN per cm^3^ of tissue. Finally, we compared PanIN density across three demographic factors to show that no significant difference in PanIN content exists as a function of patient age, sex, or location of surgical resection within this cohort (Fig 3D).

**Figure 3.**
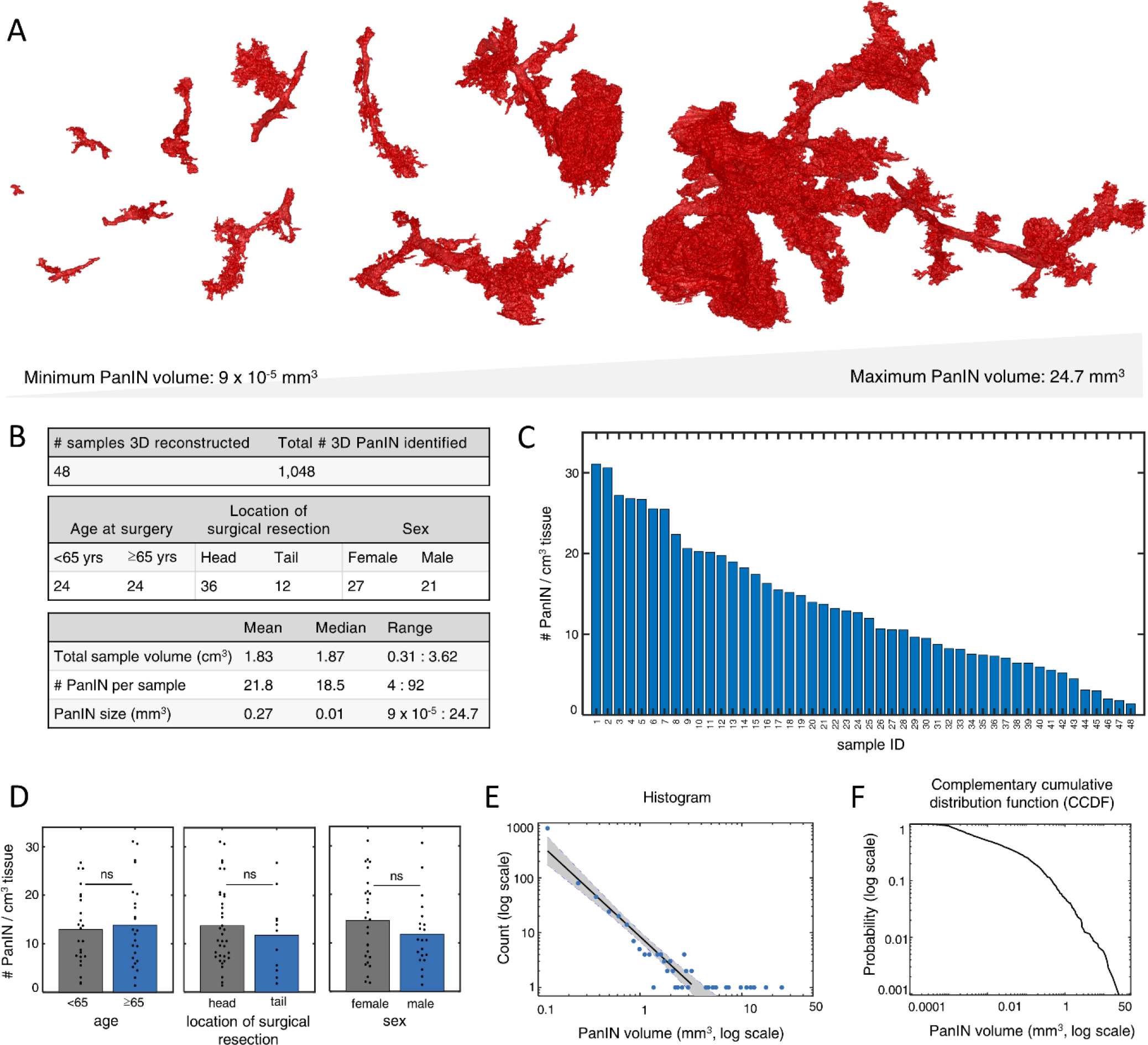
Observed PanIN sizes exhibit a broad distribution. (a) PanIN were found in a range of volumes, with a minimum PanIN volume of 9 × 10^−3^ mm^3^ and a maximum PanIN volume of 24.7 mm^3^. (b) Tables displaying total number of pancreas samples reconstructed, number of PanIN found, patient demographics, and detailed 3D sample information. (c) Bar graph displaying number of PanIN identified per cm^3^ of pancreas tissue for 48 grossly normal slabs of human tissue. Minimum of 1.4 PanIN per cm^3^ tissue and maximum of 31.1 PanIN per cm^3^ of tissue. (d) Bar graphs displaying number of Panin identified per cm3 of pancreas tissue compared across age, location of surgical resection, and sex. All nonsignificant (>0.05). (e) Histogram of PanIN volumes, plotted at logarithmic scale. (f) complementary cumulative distribution function of PanIN volumes, plotted at logarithmic scale.

The variability of PanIN size is visualized in the histogram shown in Fig. 3E. This representation of the data suggest that PanIN size is distributed according to a power law with a fitted exponent of -1.7 (correlation coefficient 0.96, with 95% confidence intervals 0.89 and 0.98), which implies that PanINs are overwhelmingly small. However, this power law cannot explain the occurrence of the largest PanIN (blue disks in lower right of Fig. 3E). A precise quantification of very large PanIN is challenging due to the limited number sampled, and this information is further concealed because histograms generally rely on binning of the data. To circumvent this problem, we instead represent the data using a complementary cumulative distribution function (CCDF), *S*(*V*), which gives the fraction of observed PanINs with a volume larger than *V* (see Fig. 3F). The precise shape of *S*(*V*) carries more information about the distribution of PanIN size than the histogram, since it does not require binning. For instance, it reveals that PanINs below *V*_*min*_ = 0.001 *mm*^3^ are rarely detected, so we disregarded data below this size in our analysis. The CCDF has a characteristic shape, which contains information about the history of the sample, i.e., when PanINs are initiated and how they grow. In the following, we test various growth models to try to explain this data.

### A power-law growth law explains size distribution qualitatively

PanIN growth is a complex, poorly understood process, which is likely affected by the pancreatic microenvironment (interactions of epithelial cells harboring somatic mutations, with stromal cells and pancreatic digestive enzymes), and an individual’s age, family history, and lifestyle. However, the comparative analysis shown in Fig. 3D suggests that age, portion of the pancreas involved (head vs tail), and sex do not significantly affect PanINs. It is thus plausible that the overall features of PanIN size distribution are less sensitive to such details and are rather shaped by general growth behavior. For instance, PanINs could grow according to their present size, proportionally to their surface area, or only along the inner lining of the pancreatic ducts in which they are, by definition, contained. These three proposed growth behaviors correspond to PanIN growth rates proportional to their volume, their surface area, and a constant, respectively. All these alternatives can be summarized by a single power-law,

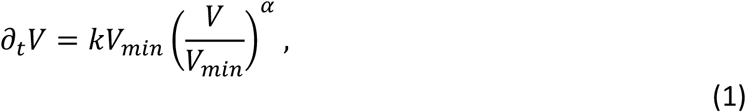

which quantifies how the PanIN volume *V* changes as a function of time *t*. Here, *V*_*min*_ is the cutoff volume, *k* quantifies the growth rate, and *α* denotes the growth exponent distinguishing different modes of growth; *α* = 1, ⅔, 0 correspond to the three alternative modes discussed above, but in principle all values of *α* are permissible. Figs. 4A and 4B visualize the strong influence of the exponent *α* on PanIN volume as a function of time.

**Figure 4.**
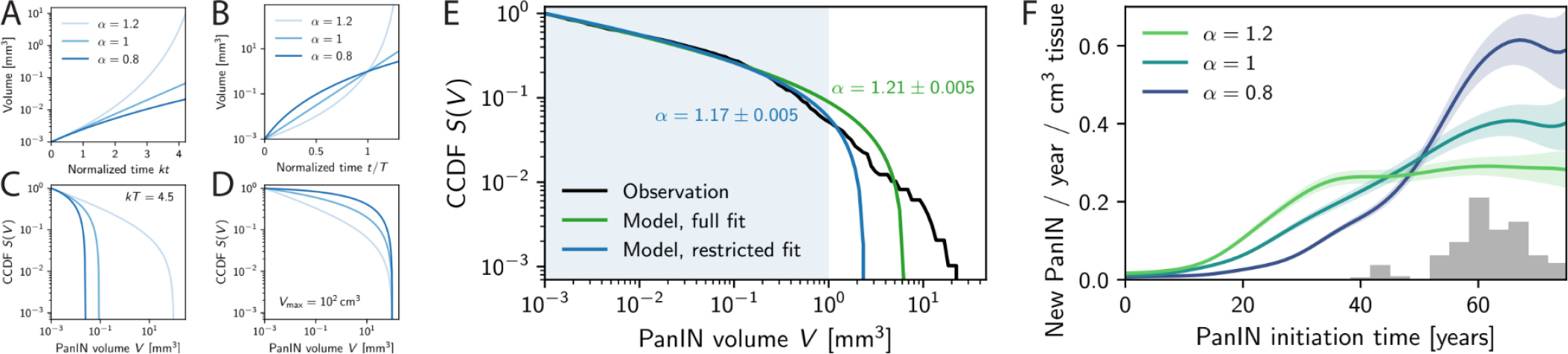
Simple growth PanIN growth model explains size distribution qualitatively. (A) PanIN volume *V* as a function of time *t* predicted by the power-law growth model given by Eq. (1) for various growth exponents *α* and identical growth rate *k*. (B) *V*(*t*) reaching the same volume at *t* = *T* for various *α*. (C) Complementary cumulative distribution function *S*(*V*) of PanIN volumes predicted by power-law growth model for a given growth rate *k* and various *α* (D) *S*(*V*) with identical maximal volume *V*_*max*_ = 100 *mm*^3^ for various *α*. (E) Comparison of observed (black line; same data as Fig. 3F) and predicted (green and blue lines) size distributions *S*(*V*). Parameters *α* and *V*_*max*_ of the power-law growth model were obtained by fitting over all volumes (green data, *χ*^2^ = 0.054) or over the indicated range (blue data, *χ*^2^ = 0.018). (F) Smoothed PanIN initiation rate density *j* as a function of age inferred using the power-law growth model and the observed PanIN sizes for various *α*. Shaded area indicates confidence interval of width 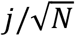, where *N* is the number of PanINs for that year. The samples ages are summarized by the gray histogram. (A-F) Additional parameters: *V*_*min*_ = 0.001 *mm*^3^

To explore suitable growth exponents *α*, we start by analyzing the simplest scenario where PanINs are initiated at a constant rate *j*, and each PanIN grows independently according to Eq.

(1). The size distribution of PanIN volumes *V* after a finite time *T* predicted by this model retains the strong dependence on *α*; see Figs. 4C and 4D. We next compare the predictions of the power-law growth model to the observed distribution *S*(*V*). Fig. 4E shows two fits of this model involving either the entire range of data (green line) or only small PanINs (blue line). This shows that the power-law growth model explains the distribution of smaller PanIN lesions reasonably well, but cannot account for the entire size distribution. This might be expected, since larger PanINs may not simply grow, but may also merge with other PanINs, which is not reflected in the current model. Nevertheless, the fit of the model to smaller PanINs suggests that PanINs grow proportionally to their volume or ever more rapidly since the model with *α* > 1 best explains the data. In contrast, the deviation of the distributions for large volumes can essentially be caused because (i) there are many more small PanINs than our simple model predicts, or (ii) there are more exceedingly large PanINs than our model predicts. Consequently, variability in PanIN initiation, but also seeding of new PanIN and merging of older PanIN lesions could explain these deviations. We will show that these scenarios are all plausible, but lead to very different dynamics, which could be discriminated experimentally.

### Growth law predicts PanIN initiation times

A core assumption of the first analysis above was that the PanIN initiation rate *j* was constant in time, whereas it is generally accepted that PanINs are more common in older individuals and that the somatic genetic events that give rise to PanINs accumulate as we age.^13,18,19^ To address this, we use Eq. (1) to predict when a PanIN measured at volume *V* must have been initiated (with a volume *V*_*min*_) relative to the age *T* of the sample. For simplicity, we use the same growth rate *k* for all PanINs, chosen minimally such that no PanINs are older than the age of the patient at the time of pancreatic resection. Taken together, this allows us to predict the initiation rate density *j* (the number of PanINs initiated in a given year per cm^3^ of pancreas tissue) as a function of time. Fig. 4F shows that a fairly constant initiation rate density requires super-exponential growth (green data), consistent with our result above. In contrast, exponential (teal data) or sub-exponential (violet data) growth requires strongly increasing initiation rates, e.g., new PanINs must appear more frequently in older samples. To get deeper insight into the connection between initiation rate *j* and the growth exponent *α*, we next discuss two concrete realizations that can cause these different behaviors.

### PanIN seeding could explain increasing initiation rates

Increased PanIN initiation rates could potentially be explained by seeding, where some neoplastic cells detach from a PanIN lesion, travel within the lumen of the duct, and initiate a new PanIN that is physically separate from the parent PanIN lesion; see Fig. 5A. Experimental evidence confirms the possibility of intraductal spread, as DNA sequencing has shown that adjacent, spatially separate PanIN sometimes harbor a similar profile of somatic mutations.^12^ To see whether this explanation is feasible, we extend the power-law growth model given by Eq. (1) to include seeding. For simplicity, we assume that the volume of a PanIN does not change when it seeds a new one, essentially assuming *V* ≫ *V*_*min*_. Seeding can then be captured by the modified initiation rate

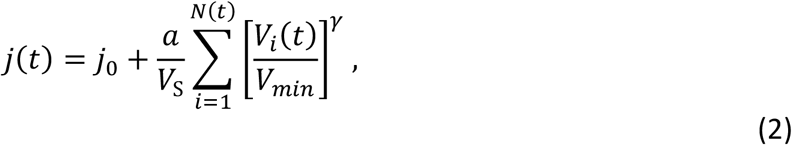

where *j*_0_ is a constant *de novo* initiation rate in the sample of volume *V*_S_, *a* quantifies the strength of seeding from each of the *N* existing PanINs of volumes {*V*_*i*_}, and *γ* is an exponent describing how the seeding depends on the size of the parent PanIN: a constant rate corresponds to *γ* = 0, whereas *γ* = 1 implies seeding proportional to the volume of the PanIN, and fractional values describe scenarios between these two extremes. Note that *j*_0_ should scale with the sample volume, whereas *a* is a rate per existing PanIN, causing an autocatalytic increase in the number of PanIN, similar to how metastasis can themselves metastasize, drastically increase the number of metastatic foci.

**Figure 5.**
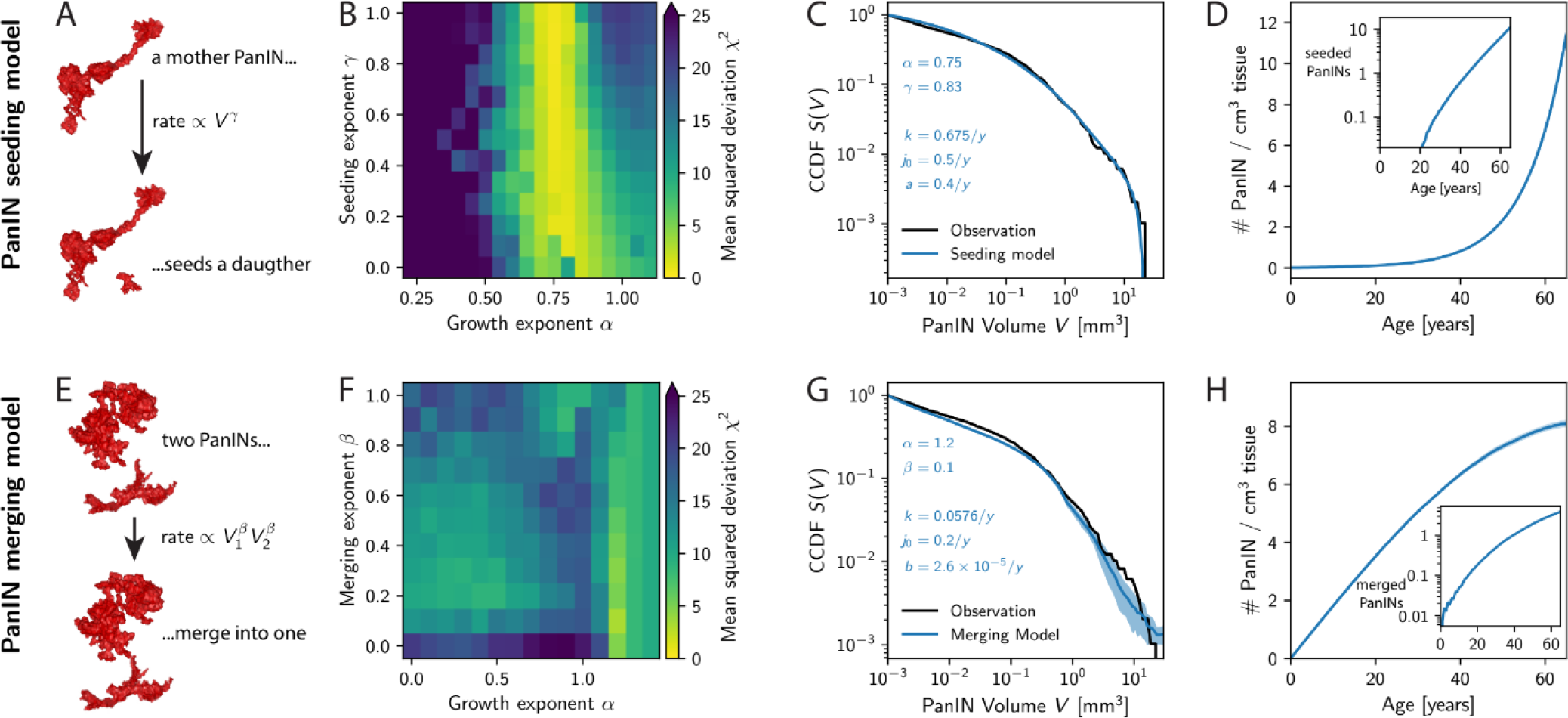
Seeding and merging models can explain observed size distribution quantitatively. These models combine the simple growth described by Eq. (1) with spontaneous seeding of daughters from older PanIN (A–D) or merging of two PanIN (E–H). (B) Mean squared deviation *χ*^2^ as a function of the growth exponent *α* and seeding exponent *γ* indicates that seeding model with *α* ≈ 0.75 can explain the observed data. (C) Comparison of the PanIN size distribution *S*(*V*) of the seeding model (blue line; *α* = 0.75, *γ* = 0.83) to the observed data (black line). The parameters in the inset refer to a sample of volume *V*_*S*_ = 100 *cm*^3^ simulated for *T* = 65 *y*. (D) Predicted PanIN count *N* as a function of age *t*. Inset shows the number of seeded PanINs as a function of *t* indicating an exponential increase. (F) *χ*^2^ as a function of *α* and the merging exponent *β* indicates that the merging model with *α* ≈ 1.2 and *β* ≈ 0.1 can explain the observed data. (G) Comparison of *S*(*V*) of the merging model (blue line; *α* = 1.2, *β* = 0.1; Shaded area indicates STD for *n* = 32 repetitions) to the observed data (black line). (H) Predicted *N* as a function of *t* suggests *N* ∼ *t*. Inset shows the number of merged PanINs as a function of age *t*.

We simulate a population of PanINs for various choices of the five parameters (*k, α, j*_0_, *a, γ*) of the PanIN seeding model to compare the resulting size distribution to the measured data. Note that two of the five parameters, namely *α* and *γ*, distinguish qualitatively different scenarios, whereas the other three parameters determine the quantitative behavior. To capture this, we analyze the model for various pairs (*α, γ*) and determine the remaining parameters using a fit to the experimental data. Using *χ*^2^ to quantify the goodness of fit, we can then judge which pair (*α, γ*) provides the best description of the experimental data. Fig. 5B shows that the seeding exponent *γ* influences *χ*^2^ only weakly, whereas the growth exponent *α* is strongly constrained by the data. Interestingly, this analysis now suggests that PanINs grow sub-exponentially (0.6 < *α* < 0.9) in contrast to the simpler model without seeding. In any case, the direct comparison of the theoretical prediction with experimental measurements shown in Fig. 5C indicates that seeding can account for the observed data quantitatively. In essence, seeding from existing PanINs leads to an exponentially increasing initiation rate *j* (see Fig. 5D), which is consistent with Fig. 4E and accounts for the many observed small PanIN lesions.

### PanIN merging could explain frequent large PanINs

A second alternative for a process that affects the size distribution are merging events where two PanINs grow so large that they touch and merge with each other within the effected duct; see Fig. 5E. Experimental evidence supports the existence of polyclonal PanIN lesions, as DNA sequencing has shown that large, highly branched PanIN lesions can contain multiple, localized containing different somatic mutations.^12^ Instead of capturing the intricate details of spatial PanIN growth, we also capture this behavior by extending of the power-law growth model given by Eq. (1). The main idea is that the probability that two PanINs meet and merge is roughly inversely proportional to sample volume *V*_*S*_ and might also depend on their individual volumes *V*_1_ and *V*_2_. We thus merge two PanINs stochastically with rate *K*(*V*_1_, *V*_2_), which we model as a power-law

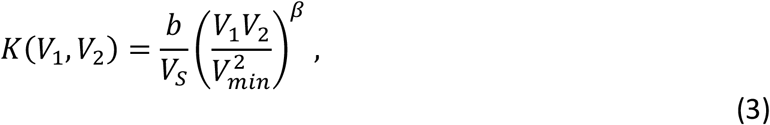

where *b* determines the merging rate, whereas *β* encodes the size-dependence: For *β* = 0, the merging rate is independent of PanIN size, whereas for instance *β* = 2/3 implies a rate that scales with the surface area of both PanINs. This merging model is similar to Smoluchowski’s coagulation model, which describes merging clusters like liquid droplets.^20,21^ For simplicity, we consider a constant rate *j*_0_ of *de novo* formation of PanINs. The model is inherently stochastic, so we simulate multiple samples and collect all PanIN volumes at the final time to compare their distribution to the experimentally measured one. Since we replace two merging PanINs by a single one with the total volume *V*_1_ + *V*_2_, this model leads to fewer but larger PanINs over time, which could explain the higher-than-expected portion of large PanINs that we observe.

The PanIN merging model has five parameters (*k, α, j*_0_, *q, β*), where again *α* and *β* distinguish qualitatively different growth scenarios, whereas *k, j*_0_, and *q* set quantitative rates. We thus again fit the rates by minimizing *χ*^2^ as a function of the parameter pair (*α, β*). Fig. 5F indicates that there is again an optimal region for these two parameters, although it is less sharply defined than in the seeding model. The best fit occurs for super-exponential growth (*α* ≈ 1.2) and a merging rate that is roughly constant (*β* ≈ 0.1), although larger merging exponents are also plausible. Fig. 5G shows that the best fit can indeed explain the observed size distribution, but there is appreciable uncertainty, particularly for the larger PanINs with worse statistics. In any case, merging of PanINs happens predominately for larger volumes, leading to even larger PanINs, implying that PanIN count decreases with time (see Fig. 5H) and the size distribution becomes skewed toward larger sizes.

### Seeding and merging model predict different PanIN counts over time

The seeding and the merging model can both explain the experimentally observed PanIN size distribution. However, the reasons are fundamentally different: The seeding model exhibits a strongly increasing initiation rate, resulting in more small PanINs than the simple power-law growth model predicts. Conversely, the merging model leads to an excess of large PanINs even for a constant initiation rate. Crucially, both models account for the deviation between the power-law growth model and the observed data that we identified in Fig. 4E. Clearly, the combination of both models could also explain the observed experimental data of PanIN sizes. However, both models make distinct predictions for the number *N* of PanINs as a function of time: The seeding model yields exponentially increasing *N* (Fig. 5D), due to the exponential increase in the initiation rate, whereas the merging model predicts even fewer PanINs than in the basic growth model due to merger events (Fig. 5H). This difference also explains why the seeding model predicts a lower growth exponent (*α* ≈ 0.75) then the merger model (*α* ≈ 1.2), which is consistent with our observations in Fig. 4D that smaller *α* coincides with strongly increasing initiation rates. Taken together, the two models could thus be distinguished, and their relative contribution quantified, if PanINs were identified in much younger samples.

## Discussion

In this work, we show that simple growth models can describe experimentally observed size distributions in human pancreatic precancer incidence and volume. We demonstrate that there are two general models of lesion growth that can lead to the experimentally measured size distribution: (1) sub-exponential lesion growth with exponentially increasing initiation rate; e.g. due to intraductal spread, and (2) exponential lesion growth with significant merging of larger lesions; e.g. fused polyclonal PanIN lesions. Both regimes fit experimentally collected genomic data – likely, a combination of the two models is true (this is studied in related fields as coagulation-fragmentation processes^22^).

Although both mechanisms lead to the same measured PanIN size distribution at their endpoints, the early dynamics of the two are very different. This is apparent in the predicted *de novo* initiation rates *j*_0_, which differ by more than two orders of magnitude (Fig. 5C and 5G), the number of lesions as a function of time (Fig. 5D and 5H), and in the lesions size distribution as a function of time (Supporting Fig. S1). The PanIN seeding model exhibits sub-exponential growth of individual PanIN lesions, but the number of PanINs grow exponentially since more PanINs can, in turn, seed more PanINs. Conversely, the PanIN merging model requires super-exponential growth of individual PanIN lesions, but the number of PanIN actually decreases over time as multiple PanIN combine into one. Since we do not observe significant differences in PanIN counts between two age groups (Fig. 3D), the merging model might explain real PanIN growth more accurately. However, reality might be best described by a combination of seeding, merging, and a time-dependent *de novo* initiation rate. More detailed data, particularly from samples from younger individuals, is needed to quantify the relative contributions of these different processes.

We note several limitations of our study. As we analyze all PanINs from all 3D samples reconstructed, our PanIN volumetric data was biased by non-fully contained lesions (PanIN that were cut at the boundaries and should thus be larger than we measure). If we were to analyze only the fully contained PanIN, we would lose all the largest lesions, shifting our distribution significantly. In the future, larger sample volumes could circumvent this problem. Because of these challenges, the numbers obtained from the model should be interpreted carefully. However, the general relations between initiation, merging, and the growth exponent would still hold. Additionally, as the volumetric PanIN data generated by CODA was limited by the resolution of the histological staining schema thickness (4 µm thick serial sections with every third section stained H&E),^12^ the resolution of the experimental data analyzed here was 12 × 12 × 12 µm^3^, which may have limited our ability to accurately measure the smallest PanIN lesions. Finally, as the pancreas samples analyzed here were collected during surgical resections for pancreatic abnormalities, the incidence and size of lesions reported here may not fully represent the general population (as most of our samples came from older individuals, and there is an association between age, pancreatic cancer, and PanIN incidence). Future work modelling the growth properties of PanIN as measured from organ donor samples and samples from younger individuals is important for correcting this bias.

Our model gives a general overview for how precancerous lesions could evolve. More detailed experimental data, e.g., based on genetic fingerprinting, would be valuable to measure seeding and merging rates directly. Similarly, more data on PanIN sizes and shapes from samples of various ages could be used to directly test different growth models of individual PanIN, e.g., whether they grow along pancreatic ducts or expand their volume in all directions (pressing outwards into the acinar lobules and inwards into the ductal luminal space), which will likely also depend on PanIN size. If such data becomes available, our model can serve as a basis for developing more detailed models which describe PanIN in the actual physical space provided by the pancreatic ducts. Moreover, our generic approach to describing lesion growth is likely transferable to other lesions types, including other common cancer precursors in the fallopian tubes or esophagus. Differences and similarities between different pre-cancerous lesions could then unveil universal principles of how cancers originate.

## Data availability statement

The data analyzed here is available from the corresponding authors upon request

## Author contributions

A.L.K., D.W., and D.Z. conceived the project. A.M.B., T.C, and R.H.H. prepared the histological specimens. A.L.K. conducted the histological image analysis. D.Z. conducted the growth simulations. L.W., P.W., D.W, and R.H.H. oversaw biological aspects of the project. A.L.K. and D.Z. wrote the first draft of the manuscript, which all authors edited and approved.

## References

1 Siegel, R. L., Miller, K. D., Wagle, N. S. & Jemal, A. Cancer statistics, 2023. CA Cancer J Clin 73, 17–48 (2023). 10.3322/caac.21763

2 Rahib, L. et al. Projecting cancer incidence and deaths to 2030: the unexpected burden of thyroid, liver, and pancreas cancers in the United States. Cancer Res 74, 2913–2921 (2014). 10.1158/0008-5472.CAN-14-0155

3 Rahib, L., Wehner, M. R., Matrisian, L. M. & Nead, K. T. Estimated Projection of US Cancer Incidence and Death to 2040. JAMA Netw Open 4, e214708 (2021). 10.1001/jamanetworkopen.2021.4708

4 Mazer, B. L. et al. Screening for pancreatic cancer has the potential to save lives, but is it practical? Expert Rev Gastroenterol Hepatol 17, 555–574 (2023). 10.1080/17474124.2023.2217354

5 Basturk, O. et al. in The American journal of surgical pathology Vol. 39 1730 (NIH Public Access, 2015).

6 Lin, J. R. et al. Multiplexed 3D atlas of state transitions and immune interaction in colorectal cancer. Cell 186, 363–381 e319 (2023). 10.1016/j.cell.2022.12.028

7 Kiemen, A. L. D., A. I.; Braxton, A.M.; He, J.; Laheru, D.; Fishman, E.K.; Chames, P.; Almagro-Perez, C.; Wu, P.W.; Wirtz, D.; Wood, L. D.; Hruban, R. H. Tissue clearing and 3D reconstruction of digitized, serially sectioned slides provide novel insights into pancreatic cancer. Med (2023).

8 Liu, J. T. C. et al. in Nature Biomedical Engineering 2021 5:3 Vol. 5 203–218 (Nature Publishing Group, 2021).

9 Richardson, D. S. & Lichtman, J. W. in Cell Vol. 162 246–257 (Cell Press, 2015).

10 Kiemen, A. L. et al. Intraparenchymal metastases as a cause for local recurrence of pancreatic cancer. Histopathology (2022). 10.1111/his.14839

11 Kiemen, A. L. et al. MRI-based Assessment of Pancreatic Fat Strongly Correlates with Histology-Based Assessment of Pancreas Composition. In press, Pancreas (2023).

12 Braxton, A.M.; Kiemen, A. L. et al. Three-dimensional genomic mapping of human pancreatic tissue reveals striking multifocality and genetic heterogeneity in precancerous lesions. biorxiv, under review (2023).

13 Carpenter, E. S. et al. Analysis of Donor Pancreata Defines the Transcriptomic Signature and Microenvironment of Early Neoplastic Lesions. Cancer Discov 13, 1324–1345 (2023). 10.1158/2159-8290.CD-23-0013

14 Fletcher, A. G., Osborne, J. M., Maini, P. K. & Gavaghan, D. J. Implementing vertex dynamics models of cell populations in biology within a consistent computational framework. Prog Biophys Mol Bio 113, 299–326 (2013). 10.1016/j.pbiomolbio.2013.09.003

15 Iwata, K., Kawasaki, K. & Shigesada, N. A dynamical model for the growth and size distribution of multiple metastatic tumors. J Theor Biol 203, 177–186 (2000). DOI 10.1006/jtbi.2000.1075

16 Kiemen, A. L. et al. CODA: quantitative 3D reconstruction of large tissues at cellular resolution. Nat Methods 19, 1490–1499 (2022). 10.1038/s41592-022-01650-9

17 Storn, R. & Price, K. Differential evolution - A simple and efficient heuristic for global optimization over continuous spaces. J Global Optim 11, 341–359 (1997). Doi 10.1023/A:1008202821328

18 Milholland, B., Auton, A., Suh, Y. & Vijg, J. Age-related somatic mutations in the cancer genome. Oncotarget 6, 24627–24635 (2015). 10.18632/oncotarget.5685

19 Risques, R. A. & Kennedy, S. R. Aging and the rise of somatic cancer-associated mutations in normal tissues. PLoS Genet 14, e1007108 (2018). 10.1371/journal.pgen.1007108

20 Cueille, S. & Sire, C. Droplet nucleation and Smoluchowski’s equation with growth and injection of particles. Phys Rev E 57, 881–900 (1998). DOI 10.1103/PhysRevE.57.881

21 von Smoluchowski, M. Three presentations on diffusion, molecular movement according to Brown and coagulation of colloid particles. Phys Z 17, 557–571 (1916).

22 Sorensen, C. M., Zhang, H. X. & Taylor, T. W. Cluster-size evolution in a coagulation-fragmentation system. Phys Rev Lett 59, 363–366 (1987). 10.1103/PhysRevLett.59.363

