## Supporting Fig. S1 for "Power-law growth models explain incidences and sizes of pancreatic cancer precursor lesions and confirm spatial genomic findings"

December 1, 2023

**2 PanIN seeding model** **4**

**3 PanIN merger model** **5**

#### 1 Simple Growth Model

This section analyzes models where lesions grow independently from each other according to the growth law

$$\partial_t V = \frac{kV^\alpha}{V_{\min}^{\alpha-1}} , \tag{1.1}$$

where $k$ quantifies the growth rate, $\alpha$ distinguishes various growth modes, and $V_{\min}$ is a reference volume.

#### 1.1 Growth of a single lesion

A single lesion, which is initiated with minimal volume $V_{\min}$ at time $t = t_0$, then exhibits the following volume $V$ as a function of time $t$,

$$V(t) = V_{\min} \cdot \begin{cases} e^{k(t-t_0)} & \text{for } \alpha = 1 \\ \left[1 + k(1-\alpha)(t-t_0)\right]^{\frac{1}{1-\alpha}} & \text{otherwise ,} \end{cases} \tag{1.2}$$

revealing that $\alpha = 1$ corresponds to exponential growth. Note that super-exponential growth ($\alpha > 1$) diverges at $t = t_{\max}$, where $t_{\max} = t_0 - [k(1-\alpha)]^{-1}$. Inverting Eq. (1.2), we can also determine the duration $\tau = t - t_0$ a lesion of volume $V$ must have been growing,

$$\tau(V) = \begin{cases} k^{-1} \log\left(\frac{V}{V_{\min}}\right) & \text{for } \alpha = 1 \\ \dfrac{1 - \left(\frac{V}{V_{\min}}\right)^{1-\alpha}}{k(\alpha - 1)} & \text{otherwise .} \end{cases} \tag{1.3}$$

This growth duration is helpful to analyze a collection of lesions.

#### 1.2 Growth of a population of lesions

We characterize a collection of lesions by the size distribution function $P(V, t)$, which gives the fraction of lesions of volume $V$ at time $t$. We define this distribution such that the integral over $V$ yields the total number $N$ of lesions at time $t$,

$$N(t) = \int_{V_{\min}}^{\infty} P(V, t)\, \mathrm{d}V \ , \tag{1.4}$$

so $P$ carries units of inverse volume. The dynamics are described by

$$\partial_t P(V, t) = j\delta(V - V_{\min}) - \partial_V \left[P(V, t)\partial_t V\right] \ , \tag{1.5}$$

where $j$ is the rate (with units of inverse time) at which new lesions of size $V_{\min}$ are introduced and the growth rate $\partial_t V$ in the second term is given by Eq. (1.1). Since Eq. (1.5) excludes lesions with volumes smaller than $V_{\min}$, the equation can also be written as

$$\partial_t P(V, t) = -\partial_V \left[P(V, t)\partial_t V\right] \qquad\qquad P(V_{\min}, t) = \frac{j}{kV_{\min}} \ , \tag{1.6}$$

where the initiation of new lesions is described by a boundary condition and $P(V, t)$ is only defined for $V > V_{\min}$.

Using the rescaled variables $v = \log(V/V_{\min})$, the growth law reads $\partial_t v = ke^{(\alpha-1)v}$ and the associated distribution $p(v, t) = V_{\min} e^v P(V_{\min} e^v, t)$ evolves according to

$$\partial_t p(v, t) = -\partial_v \big[p(v, t)\partial_t v\big] \qquad\qquad p(0, t) = \frac{j}{k} \ . \tag{1.7}$$

This form can be more suitable to evaluate expressions analytically and captures the natural spread of the lesion volume over many orders of magnitude.

#### 1.3 Size distribution for uniform nucleation

We first consider the case where lesions are continuously initiated over the time period $[0, T]$, i.e., where the initiation rate is a constant $j(t) = j_0$. The maximal volume $V_{\max}$ corresponds to the volume of the lesion initiated at $t = 0$, which reads $V_{\max} = V(T)$. This allows us to re-express the growth duration $\tau$ of a lesion of volume $V$, given by Eq. (1.3), as

$$\tau(V) = T \cdot \begin{cases} \dfrac{\log\left(\frac{V}{V_{\min}}\right)}{\log\left(\frac{V_{\max}}{V_{\min}}\right)} & \text{for } \alpha = 1 \\[2ex] \dfrac{1 - \left(\frac{V}{V_{\min}}\right)^{1-\alpha}}{1 - \left(\frac{V_{\max}}{V_{\min}}\right)^{1-\alpha}} & \text{otherwise .} \end{cases} \tag{1.8}$$

A constant rate of introduction of new lesions thus implies that $\tau \in [0, T]$, so that the cumulative distribution function $F(V)$ reads

$$F(V) = \begin{cases} \dfrac{\log\left(\frac{V}{V_{\min}}\right)}{\log\left(\frac{V_{\max}}{V_{\min}}\right)} & \text{for } \alpha = 1 \\[2ex] \dfrac{1 - \left(\frac{V}{V_{\min}}\right)^{1-\alpha}}{1 - \left(\frac{V_{\max}}{V_{\min}}\right)^{1-\alpha}} & \text{otherwise .} \end{cases} \tag{1.9}$$

The associated distribution function $P(V) = F'(V)$ of lesion sizes reads

$$P(V) = \begin{cases} \dfrac{V^{-1}}{\log\left(\frac{V_{\max}}{V_{\min}}\right)} & \text{for } \alpha = 1 \\[2ex] \dfrac{\alpha - 1}{V} \cdot \dfrac{\left(\frac{V}{V_{\min}}\right)^{1-\alpha}}{1 - \left(\frac{V_{\max}}{V_{\min}}\right)^{1-\alpha}} & \text{otherwise ,} \end{cases} \tag{1.10}$$

whereas the complementary cumulative distribution function (CCDF) or survival function $S(V) = 1 - F(V)$ reads

$$S(V) = \begin{cases} \dfrac{\log\left(\frac{V_{\max}}{V}\right)}{\log\left(\frac{V_{\max}}{V_{\min}}\right)} & \text{for } \alpha = 1 \\[2ex] \dfrac{1 - \left(\frac{V_{\max}}{V}\right)^{\alpha-1}}{1 - \left(\frac{V_{\max}}{V_{\min}}\right)^{\alpha-1}} & \text{otherwise .} \end{cases} \tag{1.11}$$

Note that the CCDF only depends on the maximal lesions size $V_{\max}$, and not on the growth rate $k$ and the growth duration $T$ individually. The CCDF is also independent of the initiation rate $j_0$, which instead affects the total number of lesions $N = j_0 T$.

##### 1.4 Predicting initiation times from size distribution

The data we want to model consists of samples of various ages $T_n$, each with several PanINs of volume $V_i^{(n)}$, where $i$ enumerates the PanINs in the $n$-th sample. We can also use Eq. (1.6) to determine the initiation rate protocols $j(t)$ that are compatible with the measured distribution at the final time points. To do this, we consider a situation where all lesions have the same growth exponent $\alpha$ (which is now an externally fixed parameter) and the same growth rate $k$. We determine the growth rate by choosing the smallest value such that the initiation times of all lesions are non-negative, i.e., such that the growth duration $\tau_i^{(n)} = \tau(V_i^{(n)})$ given by Eq. (1.3) is smaller than the sample age for all lesions, $\tau_i^{(n)} \leq T_n$. Having specified $\alpha$ and $k$, we can use Eq. (1.3) to determine the growth duration $\tau_i^{(n)}$ of each individual lesion and the associated time of initiation, $T_n - \tau_i^{(n)}$. We pool these times by years and correct for the volume of the samples and the fact that late initiation times are under-represented because we have fewer and fewer samples that grew that old; see distribution summarized in Fig. 4F of the main text. Consequently, we find the initiation rate densities $j_i$ for each year $i$. We estimate the accuracy of the values using $\sigma_i = j_i/\sqrt{N_i}$, where $N_i$ denotes the number of samples for year $i$. Finally, we smooth both $j_i$ and $\sigma_i$ with a Gaussian filter of width 5 years and plot the resulting curves $j(t) \pm \sigma(t)$ in Fig. 4F shown in the main text.

#### 2 PanIN seeding model

We model seeding by a modified initiation rate per sample volume,

$$j(t) = j_0 + \frac{a}{V_\mathrm{S}} \sum_{i=1}^{N} \left(\frac{V_i}{V_\mathrm{min}}\right)^{\gamma} , \tag{2.1}$$

where $j_0$ is the basal initiation rate, $\gamma$ determines how initiation depends on the volumes $V_i$ of individual lesions, $a$ sets the seeding rate, and $V_\mathrm{S}$ is the sample volume.

When seeding is independent of lesion size ($\gamma = 0$), the number $N$ of lesions over time is approximately governed by $\partial_t N = j_0 V_\mathrm{S} + aN$. Consequently,

$$N(t) = \frac{j_0 V_\mathrm{S}}{a} \left(e^{at} - 1\right) \tag{2.2}$$

when $N(0) = 0$, which implies $j(t) = j_0 e^{at}$. This case of an exponentially increasing initiation rate can be treated analytically and the solution to Eq. (1.6) reads

$$P(V,t) = \frac{j_0 e^{at}}{kV} \exp\left(\frac{a - a(V/V_\mathrm{min})^{1-\alpha}}{(1-\alpha)k}\right) \left(\frac{V}{V_\mathrm{min}}\right)^{1-\alpha} \tag{2.3}$$

when $\alpha \neq 1$.

The general case $\gamma \neq 0$ can only be analyzed numerically. We do this by growing and initiating lesions deterministically by evolving a collection of lesion volumes $\{V_i\}$ in time, and adding new lesions appropriately. Briefly, at each discrete time $t$, we add a

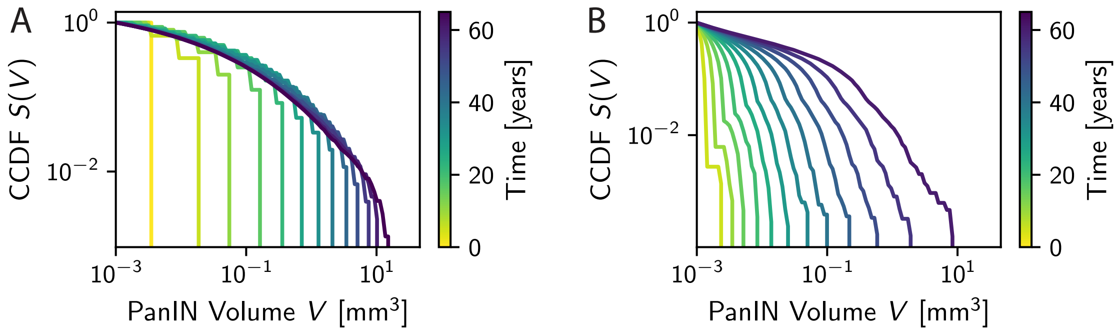

Figure S1: $S(V)$ of the seeding model (left) and the merging model (right) as a function of time (indicated by color). Model parameters are the same as in Fig. 5C and 5G of the main text, respectively.

new lesion with volume $V_{\min}$ and calculate the new waiting time $\Delta t = j^{-1}$. We then update the volume of all lesions according to

$$V(t+\Delta t) = \begin{cases} V(t)e^{k\Delta t} & \text{for } \alpha = 1 \\ \left[V(t)^{1-\alpha} + k\Delta t(1-\alpha)V_{\min}^{1-\alpha}\right]^{\frac{1}{1-\alpha}} & \text{otherwise ,} \end{cases} \tag{2.4}$$

which describes the growth described by Eq. (1.1) over a finite time interval $\Delta t$. We then update the time to $t + \Delta t$ and repeat the process until the final time $T$ is reached. This numerical implementation of the model allows us to quantify the number of lesions over time, $N(t)$, as well as the size distribution at the final time; see Fig. 5 in the main text. Fig. S1A shows the resulting complementary cumulative distribution function $S(V)$ as a function of time.

#### 3 PanIN merger model

PanIN merging is described on the mean-field level by the probability $K(V_1, V_2)$ that two particular PanINs merge,

$$K(V_1, V_2) = \frac{b}{V_S}\left(\frac{V_1 V_2}{V_{\min}^2}\right)^{\beta} , \tag{3.1}$$

where $b$ determines the rate, $V_S$ is the sample volume, and $\beta$ sets the volume-dependences. The resulting model is similar to coagulation models, e.g., when studying droplet coalescence [2]. The evolution of the distribution function $P(V, t)$ then reads

$$\begin{aligned} \partial_t P(V) &= j\delta(V - V_{\min}) - \partial_V\left[P(V)\partial_t V\right] \\ &+ \frac{1}{2}\iint K(V_1, V_2)P(V_1)P(V_2)\Big[\delta(V - V_1 - V_2) - \delta(V - V_1) - \delta(V - V_2)\Big]\mathrm{d}V_1\mathrm{d}V_2 , \end{aligned} \tag{3.2}$$

where the factor $\frac{1}{2}$ avoids double counting. This coagulation model can only be solved analytically in some special cases, and some scaling results are available [2].

To compare the predictions of the merger model to the observed data, we use a numerical solution of Eq. (3.2), based on fixed time steps $\Delta t$. During each step, we first grow all lesions, then consider potential merger events, and finally initiate new lesions if necessary. In the first part, we simply increase the volume of all lesions according to Eq. (2.4). Out of the $N_\mathrm{p} = N(N-1)/2$ total pairs of lesions, we then check $M$ random pairs $(i, j)$ and merge each of them with probability $P_{ij} = \Delta t K(V_i, V_j) N_\mathrm{p}/M$. Here, $M$ is a control parameter, which allows us to balance the computational costs against the accuracy of the simulation. We generally choose the largest value of $M$ such that $P_{ij} < 1$ for all $i, j \in 1, 2, \ldots, N$. In the final part of the time step, we initiate a new lesion if $t_\mathrm{init} < t$, where $t_\mathrm{init}$ is a pre-determined time of a next initiation event. We start with $t_\mathrm{init} = 0$ to initiate the initial lesion and then add $1/j(t)$ to $t_\mathrm{init}$ after each initiated lesion to account for time-dependent initiation rates $j(t)$. This concludes the time step, so we advance time $t$ by $\Delta t$ and repeat the procedure. This numerical implementation of the model allows us to quantify the number of lesions over time, $N(t)$, as well as the size distribution at the final time; see Fig. 5 in the main text. Fig. S1B shows the resulting complementary cumulative distribution function $S(V)$ as a function of time.

### References

[1] S. Cueille and C. Sire. Smoluchowski's equation for cluster exogenous growth. *Europhys. Lett.*, 40(3):239, nov 1997.

[2] Stéphane Cueille and Clément Sire. Droplet nucleation and smoluchowski's equation with growth and injection of particles. *Phys. Rev. E*, 57:881–900, Jan 1998.

[3] Francis Filbet and Philippe Laurençot. Numerical simulation of the smoluchowski coagulation equation. *SIAM J. Sci. Comput.*, 25(6):2004–2028, 2004.

[4] Eugenya V. Makoveeva, Dmitri V. Alexandrov, and Sergei P. Fedotov. Analysis of smoluchowski-coagulation equation with injection. *Crystals*, 12(8), 2022.
